# A distant TANGO1 family member promotes vitellogenin export from the ER in *C. elegans*

**DOI:** 10.1101/2024.07.15.603539

**Authors:** Jimmy H. Mo, Chao Zhai, Kwangsek Jung, Yan Li, Yonghong Yan, Meng-Qiu Dong, Ho Yi Mak

**Affiliations:** Division of Life Science, The Hong Kong University of Science and Technology, Hong Kong SAR, China; National Institute of Biological Sciences, Beijing, China; School of Life Sciences, Peking University, Beijing, China; Tsinghua Institute of Multidisciplinary Biomedical Research, Tsinghua University, Beijing 100084, China

**Author notes:** Current address: Department of Cell Biology and Physiology, Washington University School of Medicine in St. Louis, St. Louis, MO, USA. Current address: Touchstone Diabetes Center, University of Texas Southwestern Medical Center, Dallas, TX 75390, USA. These authors contributed equally (Chao Zhai, Kwangsek Jung, Yan Li).

## Abstract

Vitellogenin belongs to the Large Lipid Transfer Protein superfamily and is thought to share a common ancestor with human Apolipoprotein B (ApoB) for systemic lipid transport^1–5^. Vitellogenin forms part of yolk granules (also known as yolk organelles), as maternal contribution of nutrients that support the embryonic development of oviparous animals^6,7^. In *Caenorhabditis elegans*, vitellogenin proteins (VIT-1 to VIT-6) are synthesized in the hermaphrodite intestine, secreted into the pseudocoelom and internalized by oocytes^8–13^. Although a general route for inter-tissue vitellogenin transport has been described, the full mechanism that underlies its intracellular trafficking within the intestine remains obscure^9,12,13^. In humans, transport and Golgi organization protein 1 (TANGO1) and TANGO1-like (TALI) proteins generate super-sized membrane carriers to accommodate bulky ApoB-containing lipoprotein particles for their export from ER exit sites^14–17^. Transport facilitated by TANGO1 family of proteins is considered as an alternative, COPII-independent ER exit pathway^18–24^. Thus far, TANGO1 orthologs have been discovered in most metazoans, except nematodes^24,25^. Here, we report the *C. elegans* protein R148.3 (now **<u>T</u>**ra**<u>n</u>**sport and **<u>G</u>**olgi organization 1-**l**ike or TNGL-1) as a mediator of vitellogenin export from the ER. TNGL-1 depletion triggers VIT-2 accumulation in the intestinal ER lumen. TNGL-1 requires its C-terminal unstructured domain for its localization to ER exit sites. Like TANGO1, it utilizes a luminal globular domain for cargo engagement. Our findings support TNGL-1 as a distant TANGO1 family member.

## Results and discussion

### TNGL-1 shares structural similarity with human TANGO1 and TALI

We discovered TNGL-1 from a functional genomics screen for regulators of lipid accumulation in adult *C. elegans*. Analysis of the primary amino acid sequence of TNGL-1 by BLAST initially identified potential orthologs in nematodes only^26^. However, HHPred, an online server for remote protein homology detection, revealed similarity between TNGL-1 and human TANGO1 and TALI^27,28^. Indeed, AlphaFold2 predicted a shared modular structure between these proteins (Figure 1A, S1)^29^. In addition to a SH3-like globular domain near the N-terminus, TNGL-1 was predicted to harbor putative transmembrane helices, followed by a coiled-coil domain and an unstructured proline-rich domain near the C-terminus (Figure S1D-F).

**Figure 1.**
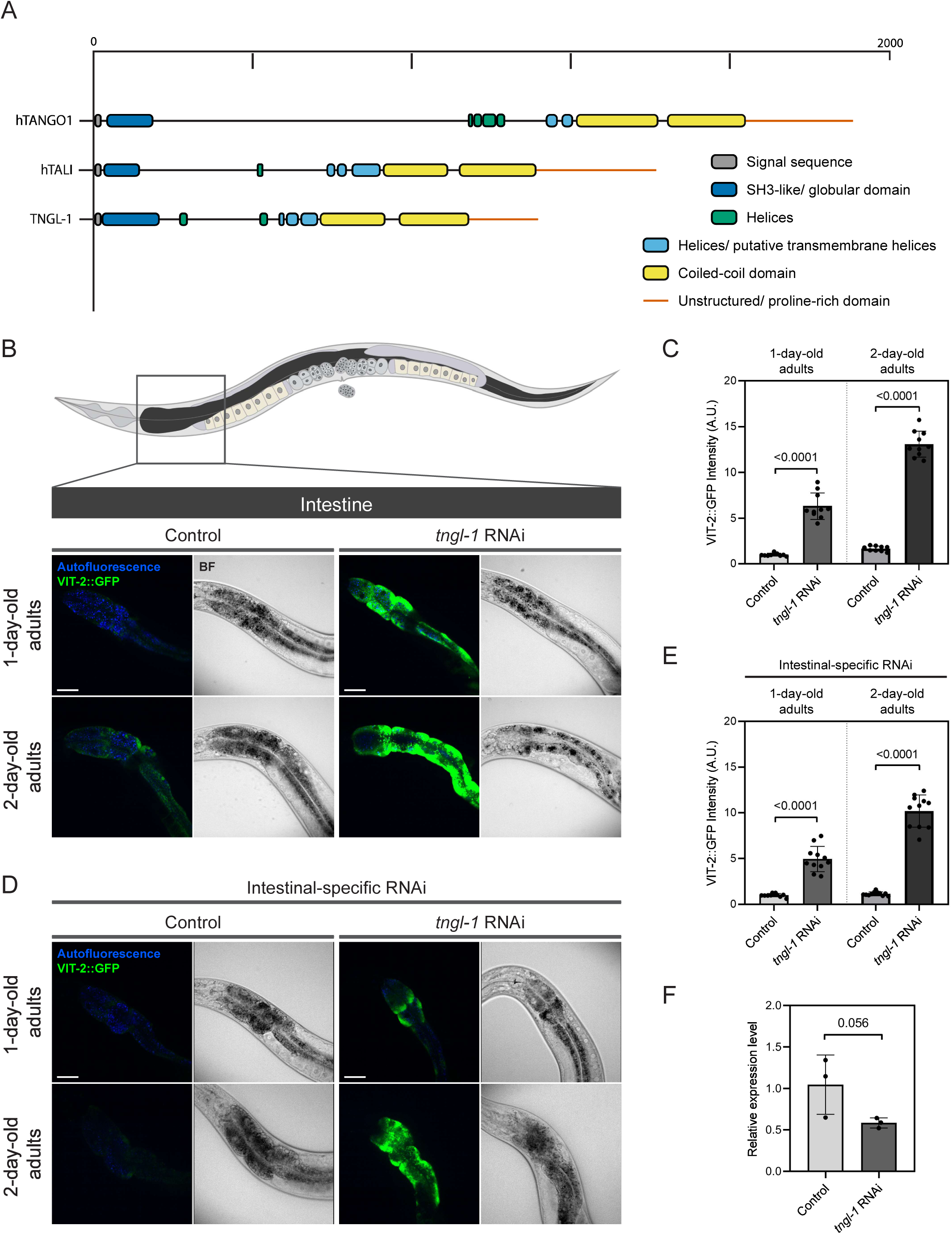
TNGL-1 deficiency causes vitellogenin accumulation in the intestine. (A) A schematic representing human TANGO1 (TANGO1, Uniprot: Q5JRA6), human TANGO1-like (TALI, Uniprot: Q96PC5), *C. elegans* TNGL-1 (Uniprot: H2KZA5). Domain boundaries are determined based on Alphafold2 predictions. (B) Top, a simplified diagram of an adult-stage *C. elegans*. The intestine is shown in dark grey. Bottom, visualization of VIT-2::GFP (*crg9070*) in 1-day-old and 2-day-old adult animals upon treatment with control or ubiquitous *tngl-1* RNAi. Images are projections of a 1 μm z stack. BF, bright field. Scale bars = 40 μm. (C) Quantification of VIT-2::GFP (*crg9070*) fluorescence intensity in the intestine. n = 10 for each group. The *p*-values are displayed (unpaired t-test). Mean ± SD is shown. (D) As in (B) but with animals subjected to intestine-specific RNAi against *tngl-1*. A single focal plane is shown. (E) As in (C), but with animals subjected to intestinal-specific RNAi. (F) Relative expression level of *tngl-1* in control or ubiquitous *tngl-1* RNAi treated animals measured by real-time PCR. *tngl-1* expression level is normalized to the average of the RNAi control group. The *p*-value is displayed (unpaired t-test). Mean ± SD from 3 biological replicates is shown.

### Partial loss of TNGL-1 impairs VIT-2 export from the ER

Human TANGO1 and TALI was previously shown to physically interact with ApoB and mediate the export of ApoB-containing lipoprotein particles^16^. To examine whether TNGL-1 has an evolutionarily conserved role, we sought to determine if TNGL-1 mediates the export of VIT-2 as a representative of vitellogenin proteins. When the mRNA level of TNGL-1 was reduced by RNA interference (RNAi) by 50%, we observed significant accumulation of the VIT-2::GFP fusion protein, expressed from the endogenous *vit-2* locus in 1-day old and 2-day old adults (Figure 1B-C, 1F). Such phenotype could be caused by an impairment in VIT-2::GFP export from the intestine or a block in its endocytosis into oocytes. To rule out the latter possibility, we conducted intestine-specific RNAi against TNGL-1 and observed a similar extent of VIT-2::GFP accumulation as before (Figure 1D-E). Our results strongly suggest that TNGL-1 promotes vitellogenin export from the intestine.

Next, we used immuno-electron microscopy to determine the localization of endogenous VIT-1 and VIT-2 at the ultra-structural level. Using gold-labeled anti-VIT-1/VIT-2 antibodies, we detected signals surrounding tubular structures in control animals (Figure 2A). In contrast, gold-labeling was detected predominantly in the lumen of membrane bound structures in TNGL-1-deficient animals. These structures are coated with granules suggesting they are likely to be the rough ER. To confirm this, we performed fluorescence microscopy on animals expressing VIT-2::GFP and an ER-luminal mCherry fluorescent protein marker (Figure 2B). Accordingly, VIT-2::GFP and mCherry colocalized in TNGL-1 deficient animals suggesting the accumulation of VIT-2 in the ER lumen (Figure 2B, panels iv-vi).

**Figure 2.**
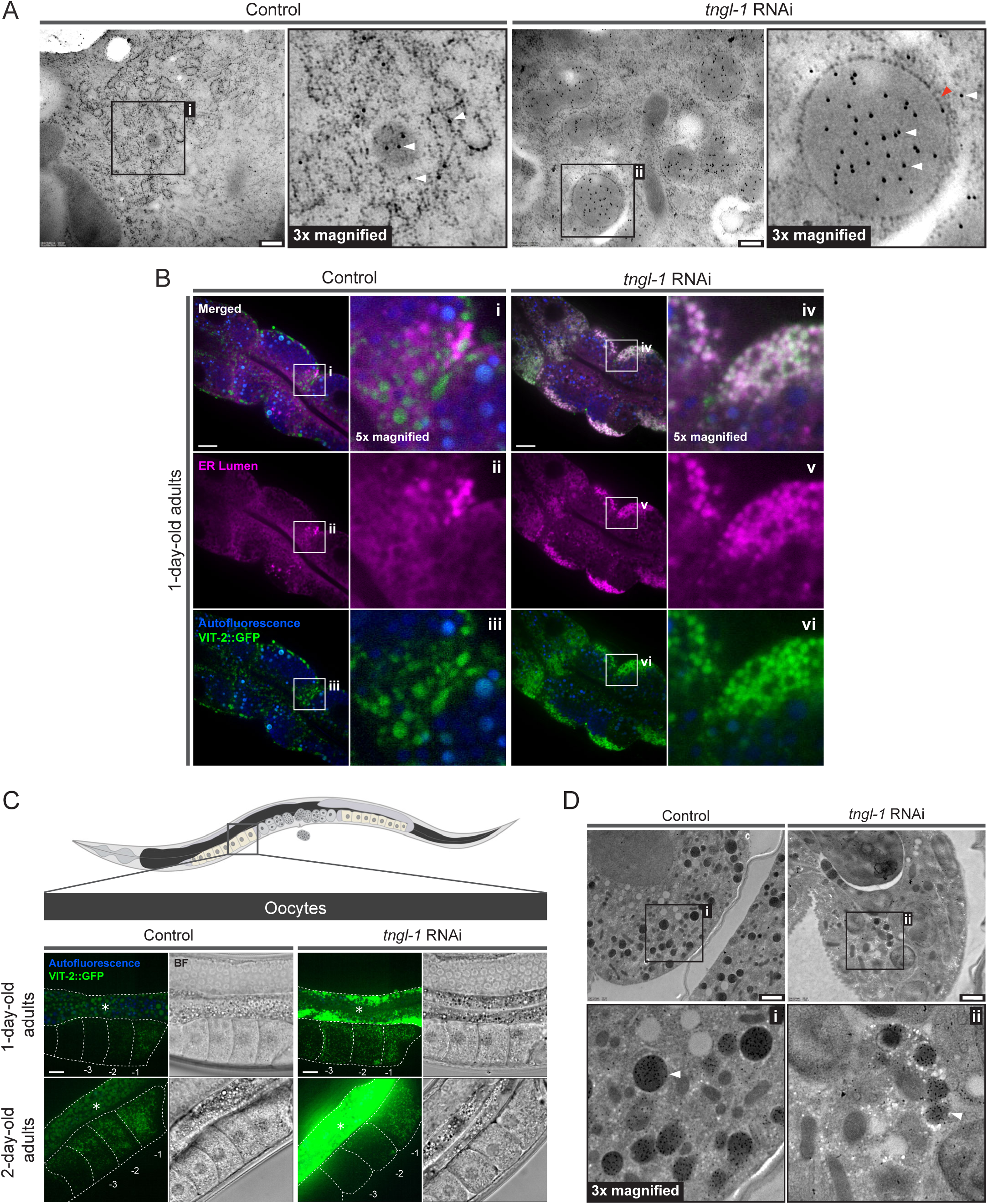
Altered vitellogenin localization in *tngl-1* deficient animals. (A) Immuno-electron microscopy images of the intestine of 1-day-old adult wild type animals. White arrows indicate anti-VIT1/2 gold particles. Red arrows indicate ribosomes. Scale bars = 200 nm. (B) Visualization of VIT-2::GFP (*crg9070*) in the intestine of 1-day-old adults upon control or *tngl-1* RNAi treatment. An ER lumen marker SEL-1(18-79)::mCherry::HDEL (*hjSi158*) was used. mCherry signal is pseudocolored magenta. Image processing parameters of vi is different from iii, with vi having reduced brightness. Images of a single focal plane centering at the second intestinal segment are shown. Scale bars = 10 μm. (C) Top, a simplified diagram of an adult-stage *C elegans*. The intestine and oocytes are in dark grey and pastel yellow respectively. Bottom, visualization of VIT-2::GFP (*crg9070*) in the oocyte of 1-day-old adults upon control or *tngl-1* RNAi treatment. Images are projections of a 1 μm z stack with the anterior, most proximal oocytes captured. Dotted lines mark the boundary between different tissues or oocytes. Asterisks mark the intestine. The oocytes closest to the spermatheca are denoted as –1, –2 and –3 respectively. BF, bright field. Scale bars = 10 μm. (D) Immuno-electron microscopy images of the embryos inside 1-day-old adult animals. White arrows indicate yolk organelles containing anti-VIT1/2 gold particles. Scale bars = 200 nm.

We noted that the partial loss of TNGL-1 function in RNAi-treated animals did not completely block the transport of VIT-2::GFP from the intestine to the oocytes, because weak green fluorescent signals could still be detected in oocytes in adult animals (Figure 2C). This was corroborated by the detection of endogenous VIT-1 and VIT-2 in yolk organelles by immuno-electron microscopy (Figure 2D). However, we could not conduct a similar analysis in animals that lack TNGL-1 completely. This was because animals homozygous for a deletion allele (*ok3525*) of *tngl-1* were inviable, as demonstrated by the inability to retrieve such animals after an outcross with wild type animals (Figure S2A). Therefore, genetic evidence suggested that TNGL-1 may have a broader function than promoting the transport of vitellogenin from the intestine to oocytes. Taken together, our results indicated that the ER exit of vitellogenin requires TNGL-1.

### TNGL-1 localizes to ER exit sites

TANGO1 and TALI are known to position at ER exit sites, presumably to promote cargo entry into the secretory pathway^16,20^. Therefore, we sought to determine if TNGL-1 was similarly found at ER exit sites. To this end, we generated a single-copy transgene that expressed a TNGL-1::mRuby fusion protein under the control of the ubiquitous *dpy-30* promoter. The TNGL-1::mRuby fusion protein was functional because its expression rescued the *tngl-1* deletion mutant animals and supported their development into reproductive adults (Figure S2B). In animals that co-expressed TNGL-1::mRuby and the SEC-16A.1::GFP ER exit site marker, we detected co-localization of green and red fluorescence signals in punctate structures in the intestine (Figure 3A). Furthermore, it was shown that TANGO1 required its C-terminal region for correct targeting to ER exit sites^20^. In particular, the proline tripeptide motifs (PPP motifs) found in this region increases its binding affinity to Sec23, an ER exit site component^30^. Therefore, we fused mRuby to a series of mutant TNGL-1 proteins that harbored progressive C-terminal truncations. We used the AlphaFold2 prediction to guide the design of the truncation mutants, to minimize gross structural disruption by placing the deletion boundaries within predicted unstructured loop regions. We observed that the deletion of both PPP motifs did not disrupt the localization of TNGL-1 to ER exit sites (Figure 3B-C, S3A-D). However, fewer TNGL-1::mRuby puncta co-localized with the ER exit site marker SEC-16A.1::GFP, when the unstructured proline-rich domain of TNGL-1 was also deleted (Figure 3D, S3E). Further deletion of the coiled-coil domain completely abrogated the localization of TNGL-1 to ER exit sites (Figure 3E, S3F). In agreement to the above, the fusion proteins that failed to localize to ER exit sites were dispersed within the ER network as revealed by the ER-membrane marker ACS-22::GFP^31^ (Figure S3E-F). Our results indicated that TNGL-1 and TANGO1 relied on different C-terminal domains for their targeting to ER exit sites.

**Figure 3.**
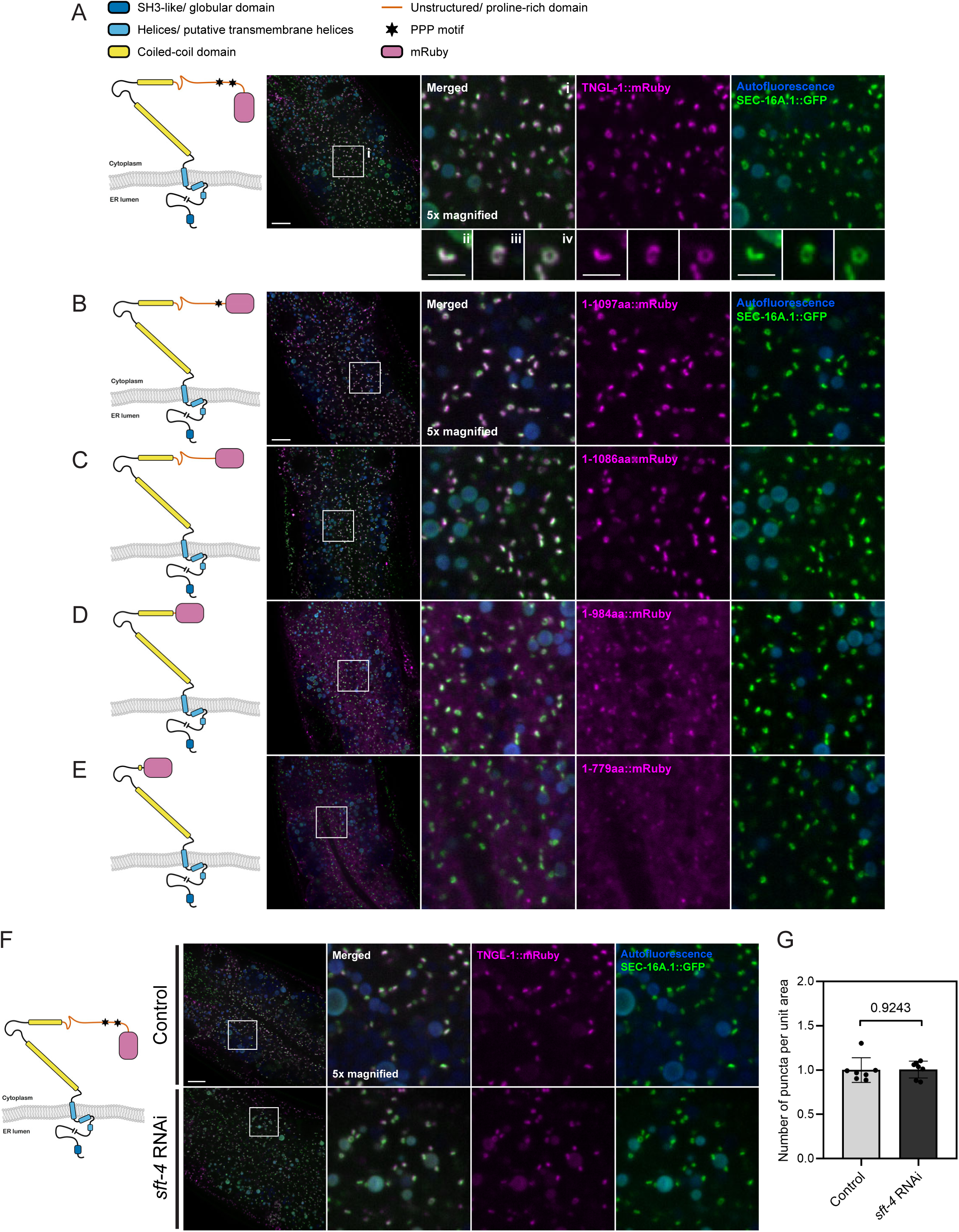
TNGL-1 localizes to the ER exit site. (A) Left, a simplified representation of the TNGL-1::mRuby fusion protein structure. Not drawn to scale. Right, visualization of TNGL-1::mRuby (*hjSi636*) and SEC-16A.1::GFP (*hj256*) in the intestine of 1-day-old adults. mRuby signal is pseudocolored magenta. Representative images of TNGL-1::mRuby signal pattern were 10x magnified and displayed at the bottom. Images are projections of a 1 μm z stack. Top scale bar = 10 μm, bottom scale bars = 2 μm. (B-E) as in (A) but represent versions of TNGL-1::mRuby fusion protein with the C-terminus progressively truncated. Boxed regions were magnified 5x and displayed on the right. Scale bar = 10 μm. (F) as in (A) but with animals subjected to control or *sft-4* RNAi. Scale bar = 10 μm. (G) Quantification of the number of SEC-16A.1::GFP positive puncta per unit area in control or *sft-4* RNAi treated animals. n = 7 for each group. The *p*-values are displayed (unpaired t-test). Mean ± SD is shown.

The *C. elegans* SFT-4 and its mammalian ortholog SURF4 promote the ER export of vitellogenins and lipoproteins, respectively, as cargo receptors^32,33^. It is unknown if SFT-4 and SURF4 coordinate with TANGO1 family of proteins in this process. We confirmed that the depletion of *sft-4* by RNAi caused VIT-2::GFP retention in the intestine (Figure S4A). However, the localization of TNGL-1::mRuby to ER exit sites was not severely disrupted in SFT-4 deficient animals (Figure 3F-G). On the other hand, TNGL-1 depletion caused an aggregation of SEC-16A.1::GFP and mRuby::SEC-24.2 fusion proteins (Figure S4B). This morphological change resembles that when TANGO1 was depleted in mammalian cells^20^. Furthermore, the number of ER exit sites decreased upon *tngl-1* RNAi (Figure S4C). The different effects on ER exit site morphology upon their depletion highlight overlapping but distinct roles of SFT-4 and TNGL-1 in ER cargo export.

### TNGL-1 interacts with VIT-2 using its N-terminal SH3-like globular domain

TANGO1 is incapable of procollagen binding in the absence of its SH3-like domain^20^. In addition, mice harboring a mutation in the SH3-like domain of TALI has reduced blood cholesterol and triglycerides level^15^. These observations point to the SH3-like domain being responsible for cargo engagement. Therefore, we hypothesized that the N-terminal SH3-like globular domain of TNGL-1 has an equivalent role. To test this, we first performed immunoprecipitation experiments using lysates of worms that co-expressed TNGL-1::mRuby::3xHA and VIT-2::GFP fusion proteins (Figure 4A). We found that VIT-2::GFP co-immunoprecipitated with TNGL-1::mRuby::3xHA reproducibly (Figure 4B, lanes 9 and 12). As a negative control, the ER membrane resident ACS-22::GFP^31^ did not co-immunoprecipitate with TNGL-1::mRuby::3xHA (Figure 4B, lanes 8 and 11). Therefore, TNGL-1 and VIT-2 specifically associate with each other. Next, we designed an *in vitro* binding assay to determine if the SH3-like domain alone is sufficient for VIT-2::GFP binding. We purified bacterially expressed TNGL-1 SH3-like domain as GST-fusion proteins and incubated them with whole worm lysate containing VIT-2::GFP (Figure 4C and 4D, lane 7). In parallel, we used GST alone as a negative control (Figure 4C and 4D, lane 6). The GST-TNGL-1 SH3-like domain, but not GST alone, was able to bind VIT-2::GFP (Figure 4D, lanes 4 and 5). We noticed only a small portion of VIT-2::GFP was captured in both our *in vivo* and *in vitro* binding assay when whole-worm lysates were used (Figure 4B and D). Vitellogenin is known to undergo post-translational modification following their ER export^34,35^. Moreover, *C. elegans* vitellogenin is found as complexes in embryos^36^. It is plausible that the TNGL-1 SH3-like domain binds only transiently to VIT-2 when they are at the intestinal ER. The majority of VIT-2 disengages with TNGL-1 during subsequent trafficking steps. Taken together, our results demonstrate physical interaction between TNGL-1 with VIT-2, supporting the notion that TNGL-1 is a TANGO1-family member.

**Figure 4.**
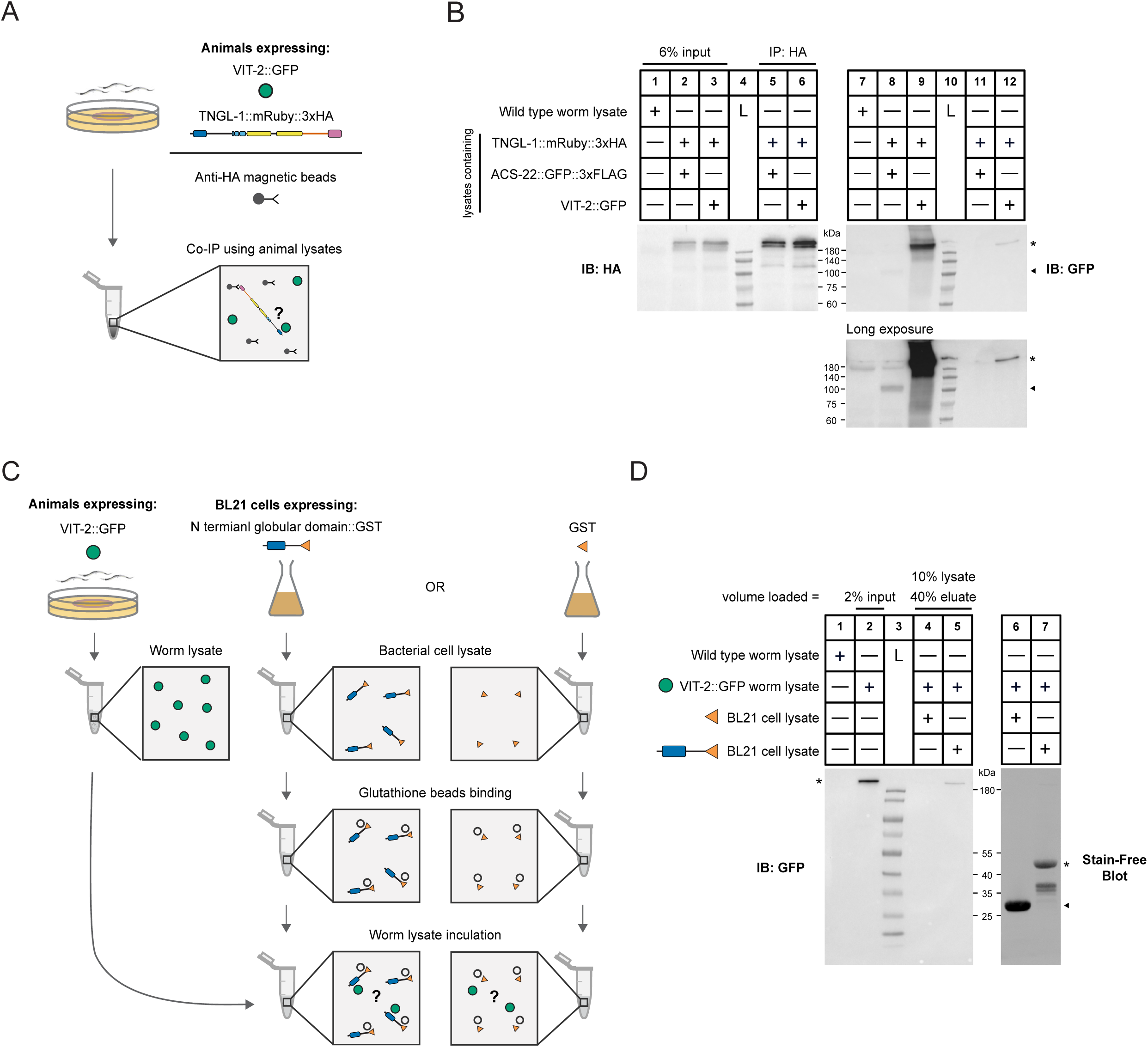
TNGL-1 N-terminal globular domain interacts with VIT-2::GFP. (A) Schematic depicting the experimental workflow of the co-immunoprecipitation experiment using whole worm lysate. (B) Co-immunoprecipitation of TNGL-1::mRuby::3xHA (*hjSi636*) and VIT-2::GFP (*crg9070*) (asterisk). Lysate made from animals expressing TNGL-1::mRuby::3xHA (*hjSi636*) and ACS-22::GFP::3xFLAG (*hjSi29*) (arrowhead) was used as a negative control. The samples loaded for the blot on the left and right are the same, but the blots are incubated with different antibodies. (C) Schematic representing the experimental workflow of the *in vitro* binding assay. (D) Binding of GST::TNGL-1 N-terminal globular domain (A23-G198) (asterisk) with VIT-2::GFP. GST protein (arrowhead) was used as a negative control. Lanes 6 and 7 correspond to lanes 4 and 5, respectively, but with GST proteins visualized using stain-free imaging. L, ladder. IB, immunoblot. IP, immunoprecipitation.

### Conclusions

Cargo export from the ER is a fundamental cellular process. The evolutionarily conserved TANGO1 family of proteins have been identified as export facilitators for cargos that cannot fit into COPII vesicles. Paradoxically, no *C. elegans* family members have been described^24,25^. TNGL-1 was previously proposed to modulate lipid metabolism and longevity but is thought to act as a signaling molecule^37^. In this study, using live imaging, electron microscopy and biochemical assays, we provided strong evidence to support TNGL-1 as a distant TANGO1 family member that promotes vitellogenin export in *C. elegans*. Moreover, we defined TNGL-1 domains that are necessary for its correct localization to the ER exit site and for vitellogenin binding.

The identification of *C. elegans* TNGL-1 implies that TANGO1 protein family-dependent export of systemic lipid carrier precursors from the ER is broadly conserved. However, the mechanism by which these lipid carrier precursors interact with lipids may be distinct. Lipoproteins in mammals arise from the ER containing a neutral lipid core delimited by a monolayer of phospholipids^4,38^. During their synthesis, apolipoproteins such as ApoB associate with their surface^39–41^. In contrast, *C. elegans* vitellogenin has been observed within membrane bound vesicles during their transit to stations of the secretory pathway^42^. How do structurally similar TANGO1 and TNGL-1 engage distinct lipid carriers? TANGO1 is known to mediate the export of discrete cargoes and such broad activity is thought to be enabled by adaptors^24,25^. One possibility is that these accessory components are also required in *C. elegans*, but the adaptor that participates in the ER export of vitellogenin and ApoB-containing lipoprotein particles are divergent. Future study will explore how cargo selection for ER export mediated by TANGO1 family of proteins have evolved.

In addition to lipoproteins, TANGO1 is also required for the secretion of procollagen^20^. Interestingly, TMEM-131 and TMEM-39 have recently been reported as conserved mediators of collagen export in *C. elegans*^43,44^. With the identification of TNGL-1, further study is necessary to clarify if TNGL-1 acts together or independently with TMEM-131 and TMEM-39. Beyond bulky cargos, *Drosophila* Tango1 appears to contribute to general protein secretion^24,45^. Therefore, TANGO1 family of proteins may have diverged in separate lineages to facilitate the export of different range of cargos. Intriguingly, TNGL-1 deletion in *C. elegans* renders the animal inviable. This could be caused by a failure to transport specific essential cargoes or an impediment to protein secretion in general. Studying the interactome of TNGL-1 will distinguish between these possibilities and further shed light on how protein export machineries evolved in different branches of metazoans.

## STAR Methods

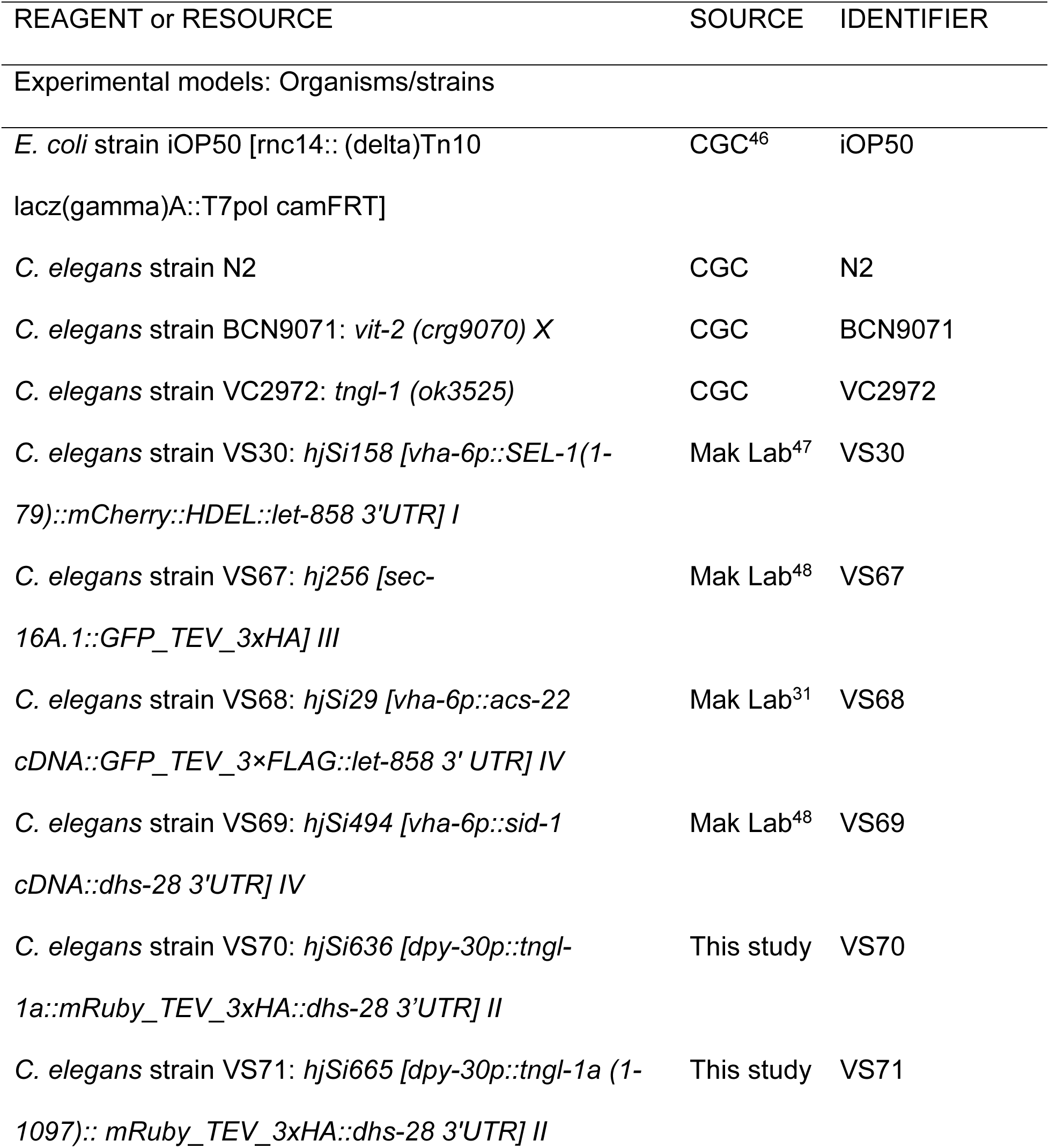
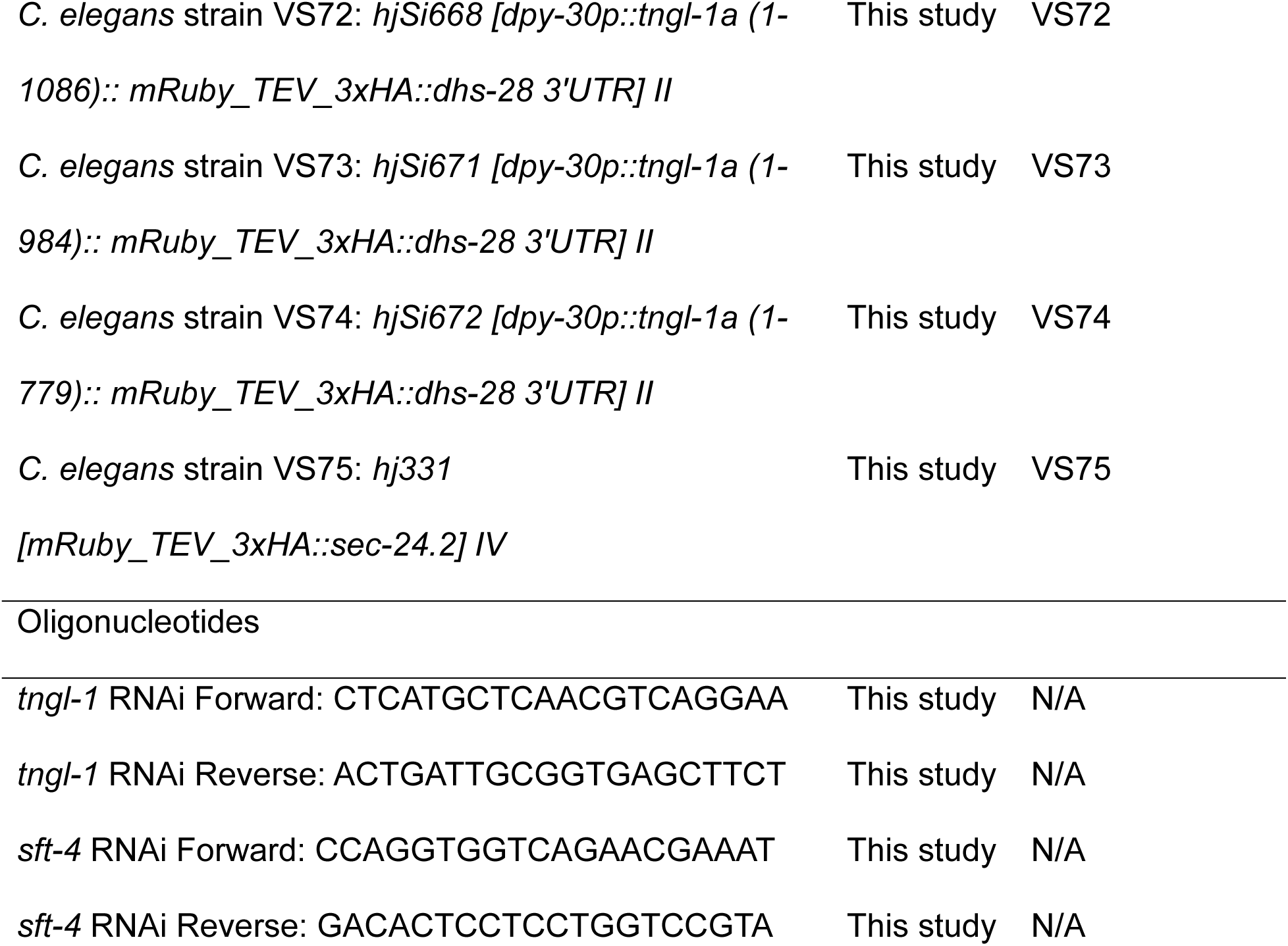
Key resources table.

## Resource availability

### Lead contact

Further information and requests for resources and reagents should be directed to the lead contact, Ho Yi Mak.

### Materials availability

*C. elegans* strains generated in this study are available from the lead contact upon request.

### Data and code availability

All data reported in this paper will be shared by the lead contact upon request. No code is associated with this paper.

## Experimental models and study participant details

### Strain maintenance and generation

*C. elegans* were propagated on NGM plates seeded with *E. coli* strain OP50 at 20°C^49^. A complete list of strains used in this study can be found in the key resources table. Transgenes were generated using CRISPR-Cas9 based techniques as described previously^50^. Knock-in alleles were generated by injecting plasmid DNA expressing Cas9 and sgRNA designed using CHOPCHOP v3^51^. Single-copy transgenes were generated as the above but sgRNA targeting the MosI sites of MosSCI strains were used^52^. All generated *C. elegans* strains were outcrossed with N2 at least twice and the inserted sequences were verified using Sanger sequencing.

## Method details

### RNA Interference-based Knockdown in *C. elegans*

RNA interference (RNAi) experiments were performed based on published methods^46^. RNAi constructs for *tngl-1* and *sft-4* were transformed into *E. coli* iOP50. The target sequences were amplified from cDNA using primers detailed in the key resources table. Fresh overnight liquid cultures of OP50 RNAi clones were seeded on NGM plates containing 0.4mM IPTG and 100µg/mL Ampicillin. The seeded plates were stored in the dark and incubated at room temperature overnight. Unless otherwise specified, for *tngl-1* RNAi, 10 L4 P0 animals were transferred onto RNAi plates. A day later when these animals became 1-day-old adults, they were transferred onto a new batch of RNAi plates for 2 hours of egg-laying and subsequently removed. The progenies were assayed. For *sft-4* RNAi, synchronized L1 larvae were seeded onto RNAi plates. The same animals were later assayed.

### Fluorescence Imaging

Imaging samples were prepared by mounting live *C. elegans* on fresh 8% agarose pads with 0.5 mM levamisole (Sigma) in 1×PBS on microscope glass slides. 1-day-old adult worms were examined by a spinning disk confocal microscope (AxioObeserver Z1, Carl Zeiss) using either a 20x, numerical aperture (NA) 0.8, or 63x, numerical aperture (NA) 1.4 oil Alpha-Plan-Apochromat objectives. Image stacks were acquired by a Neo sCMOS camera (Andor) controlled by the iQ3 software (Andor). For GFP, a 488 nm laser was used for excitation, and signals were collected with a 500–550 nm emission filter. For mRuby, a 561 nm laser was used for excitation, and signals were collected with a 580.5–653.5 nm emission filter. For autofluorescence from lysosome-related organelles, a 405 nm laser was used for excitation, and signals were collected with a 417–477 nm emission filter. Images were exported to Imaris 8 (Bitplane) for processing. Signal quantification and puncta analysis were performed using ImageJ.

### Immuno-electron Microscopy

Immuno-electron microscopy was performed as previously described^42^. RNAi worm samples were prepared as mentioned above but 25 P0 animals were transferred onto RNAi plates for egg laying and removed after 4 hours. As the progenies became 1-day-old adults, they were collected for sample preparation.

### Real-time qPCR

RNAi worm samples were prepared as mentioned above but 50 P0 worms were transferred onto RNAi plates for 2 hours egg-laying to yield about 400 progenies. As the progenies became 1-day-old adult worms, they were harvested for RNA extraction. Total RNA was extracted with a Direct-zol^TM^ RNA MiniPrep kit (Zymo). 400ng of total RNA was reverse transcribed with a Transcriptor cDNA synthesis kit (Roche). Real-time qPCR was conducted using a Roche LightCycler system with SYBR Green Master Mix (Roche). The Delta-delta CT method was used for analyzing the raw CT values. The following primers were used for RT-qPCR: *tngl-1*, 5’-GGAAGAGCTCACCACACCTC-3’ and 5’-ACTCGCAGCTACAAACGTCA-3’; *rpl-32* (internal standard), 5’-AGGGAATTGATAACCGTGTCCGCA-3’ and 5’-TGTAGGACTGCATGAGGAGCATGT-3’.

### Immunoprecipitation

Worm samples were prepared from about 8000 synchronized 1-day-old adult animals. These animals were harvested and lysed by mechanical grinding followed by incubation at 4°C for 2 hours with end-over-end rotation in 1X MAPK Lysis Buffer (10mM Tris-HCl pH8.0, 50mM NaCl, 1mM EDTA, 0.5% IGEPAL CA-630) with 2X protease inhibitor (Complete, Roche). The soluble fraction of the lysates was then incubated with Pierce Anti-HA Magnetic Beads (Thermo Fisher) at 4°C for 2 hours with end-over-end rotation. The beads were washed 3 times using 1X MAPK Lysis Buffer with 1X protease inhibitors (Complete, Roche). The bound proteins were then eluted by boiling at 70°C for 10 minutes in 1×SDS loading buffer prior to SDS-PAGE. For Western blotting, anti-GFP (A-11122, Thermo Fisher) and anti-HA (11867423, Roche) antibodies were used at dilutions of 1:2500 and 1:2000 for detecting GFP-tagged and HA-tagged proteins, respectively.

### In Vitro Binding Assay

Worm samples were prepared from about 4000 synchronized 1-day-old adult animals maintained on NGM plates seeded with standard *E. coli* OP50 food. Worm lysates are prepared as described in the co-immunoprecipitation assay section. Expression of GST fusion proteins in bacteria was conducted as follows. Vectors expressing TNGL-1 A23-G198::GST fusion and GST tag alone were transformed into BL21 (DE3) cells. Overnight cultures incubated at 37°C were diluted 1:20 in LB medium containing 100 μg/mL Ampicillin. Refreshed cultures were incubated at 37°C until OD600 reached 0.6-0.7. Expression was induced using 0.05mM isopropyl β-D-1-thiogalactopyranoside (IPTG) at 16°C overnight. Bacterial cells were collected by centrifugation and resuspended in 1X MAPK Lysis Buffer with 1X protease inhibitor (Complete, Roche). They were subsequently lysed by sonication. The soluble fraction of the bacterial lysate was used to incubate with Glutathione Sepharose™ 4B beads (17075605, GE Healthcare) at 4°C for 1 hour with end-over-end rotation and washed 3 times with 1X MAPK Lysis Buffer with 1X protease inhibitor (Complete, Roche). Worm lysates prepared from the above were then equally divided into tubes containing glutathione beads coated with respective proteins for 2 hours incubation at 4°C with end-over-end rotation. After that, the glutathione beads were washed 3 times with 1X MAPK Lysis Buffer with 1X protease inhibitor (Complete, Roche). Proteins bound were eluted using 1xSDS loading buffer and boiled at 70°C for 10 minutes before SDS-PAGE using a stain-free gel (1610183, Bio-Rad). After protein transfer from the gel to the membrane, proteins are visualized using stain-free imaging on a Chemidoc imaging system (Bio-Rad). VIT-2::GFP proteins were visualized by western blotting using an anti-GFP antibody (A-11122, Thermo Fisher) at a dilution of 1:2500.

### Statistical analyses

All statistical analyses were performed using GraphPad Prism software. Statistical tests are detailed in the Figure Legends. Error bars represent standard deviation and ‘n’ refers to the number of animals tested in a single experiment. *p-value* for each comparison is displayed.

## Acknowledgments

We thank Ziyun Ye for assisting with *in vitro* binding experiments and Siwei Huang for RT-qPCR experiments. We thank Vivek Malhotra for advice, David Banfield and all members of the Mak and Dong labs for helpful discussions. We also thank the Biosciences Central Research Facility (Clear Water Bay), HKUST for technical support. This work was supported by RGC GRF 16101820.

## Author contributions

J.H.M. was responsible for Fig. 1A-C, E-F, 2B-C, 3, 4, S1, S2, S3, S4. Z.C. was responsible for Fig. 2A, D. K.J. was responsible for Fig. 1D, 3B-E, S3B-F. Y.L. was responsible for the functional genomic screen, preliminary experiments for Fig. 1B, 2C, S2A. Y.Y. was responsible for curating a gene set for the functional genomic screen. J.H.M. and H.Y.M. wrote the manuscript. H.Y.M. and M.Q.D. supervised the project.

## Figure Legends

**Figure S1.**
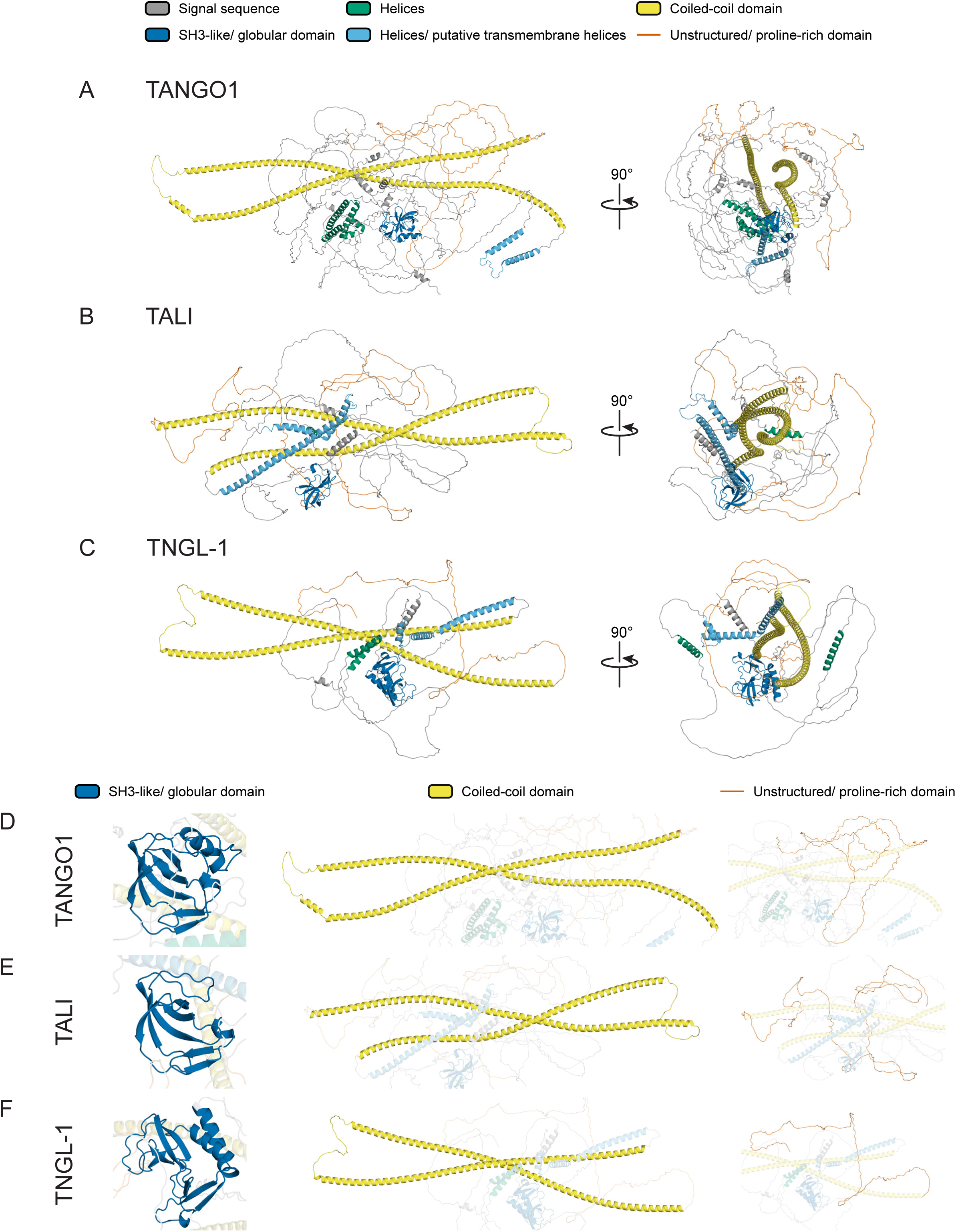
Alphafold2 structural predictions of. (A) human TANGO1 (Uniprot: Q5JRA6), (B) human TALI (Uniprot: Q96PC5) and (C) *C. elegans* TNGL-1 (Uniprot: H2KZA5). (D) Predicted structure of TANGO1 SH3-like N-terminal globular, coiled-coil and C-terminal unstructured proline-rich domains. (E, F) As in (D), but domains of TALI and TNGL-1 are shown respectively. Visualizations of the predicted structures were made using PyMOL (The PyMOL Molecular Graphics System, version 2.0, Schrödinger, LLC.).

**Figure S2.**
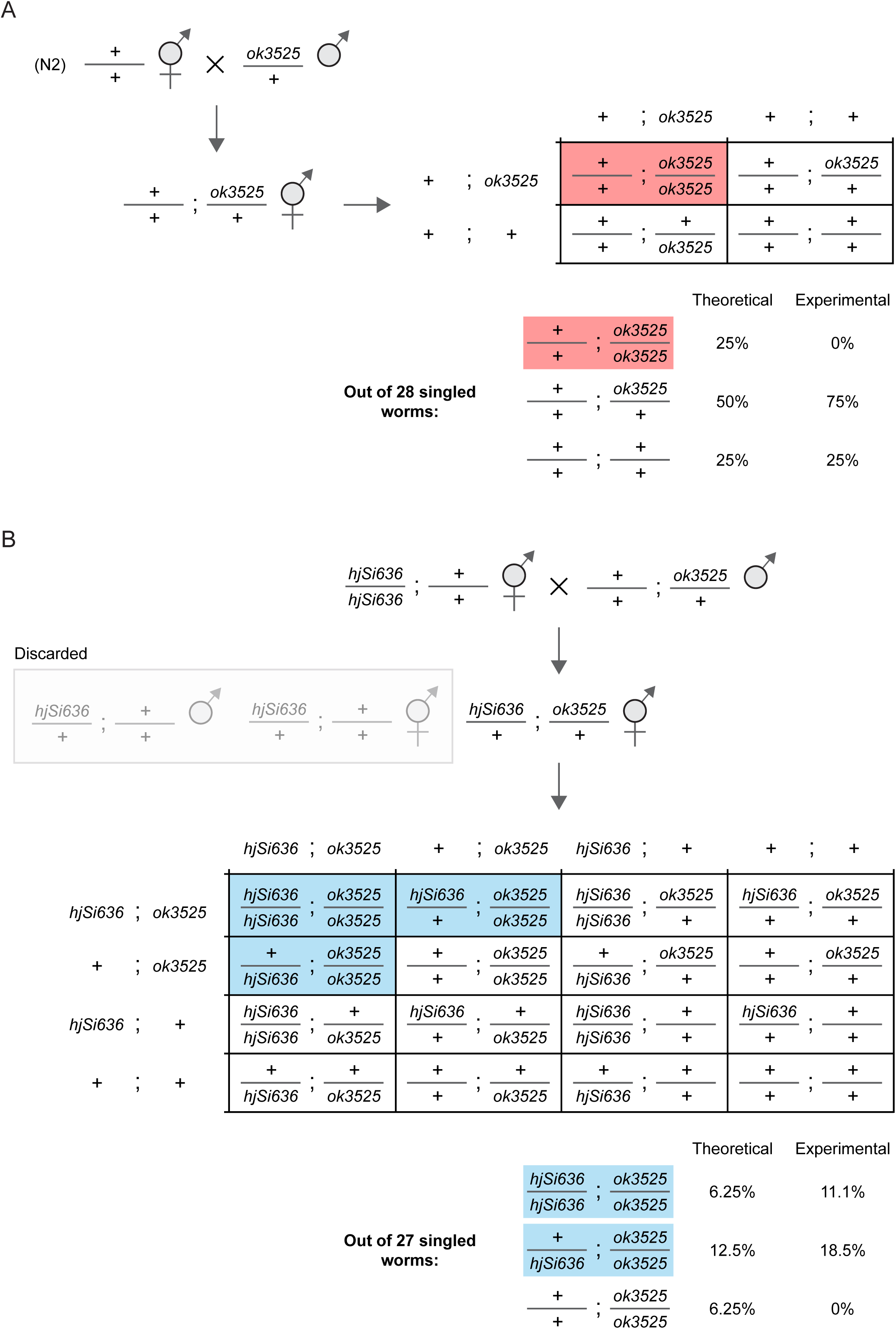
Genetic crosses indicate TNGL-1::mRuby (*hjSi636*) is functional. (A) A genetic cross between wild type N2 and *tngl-1 (ok3525)* knockout animals. The genotype that is expected to be embryonic lethal is marked by a red box. (B) A genetic cross of *hjSi636* and *tngl-1 (ok3525)* knockout animals. Genotypes suggesting rescue are marked by blue boxes.

**Figure S3.**
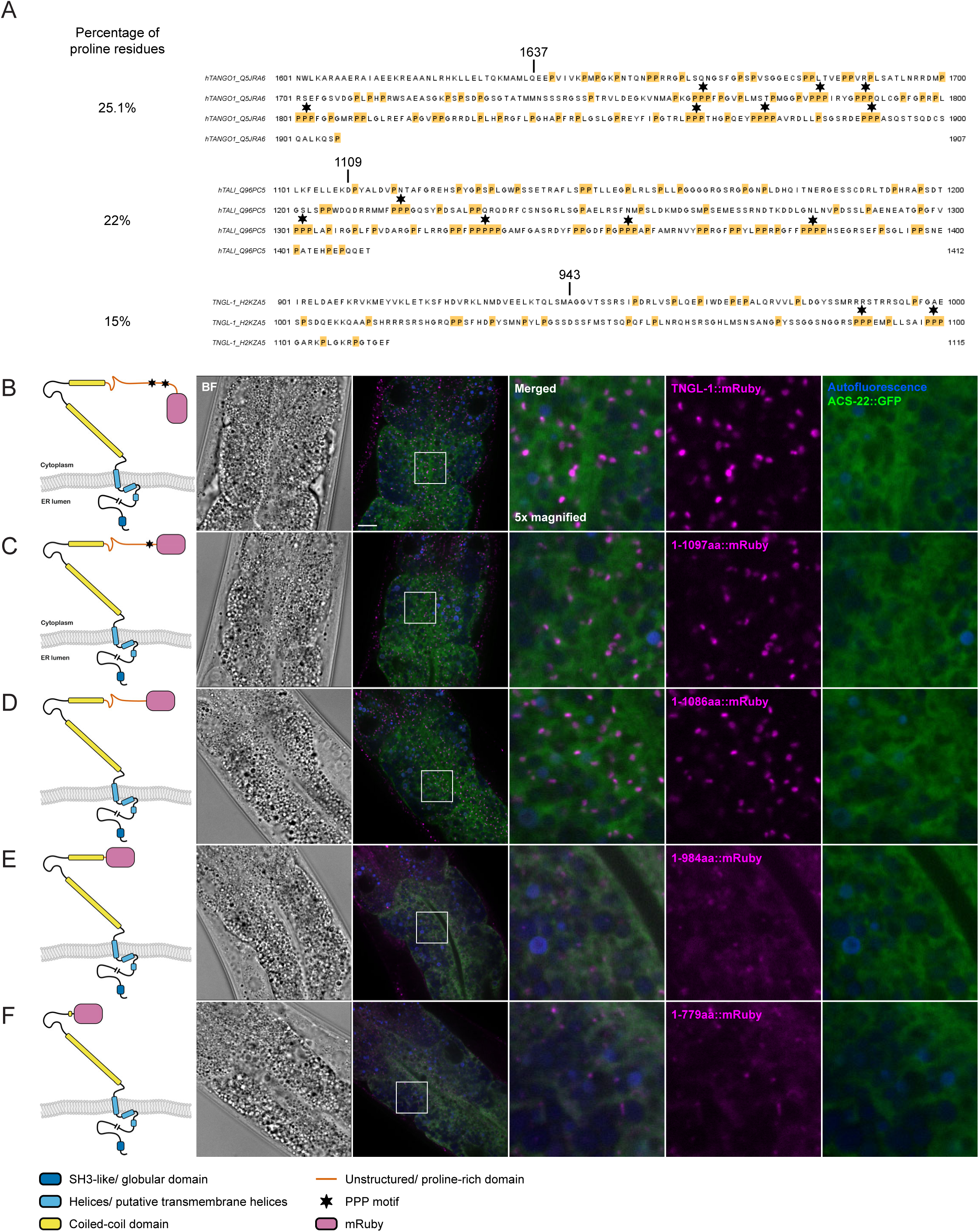
TNGL-1 C-terminal unstructured region, but not the proline tripeptide motifs, is required for proper localization of TNGL-1 to ER exit sites. (A) Protein sequences of C-terminal unstructured region for hTANGO1 (starting at residue 1673), hTALI (starting at residue 1109), TNGL-1 (starting at residue 943). Asterisks mark the position of proline tripeptide motifs (PPP motifs). The percentage of proline residues in the C-terminal unstructured region of each protein is summarized on the left. (B) Left, a simplified representation of the TNGL-1::mRuby fusion protein structure. Not drawn to scale. Right, visualization of TNGL-1::mRuby (*hjSi636*) and ACS-22::GFP::3xFLAG (*hjSi29*) in the intestine of 1-day-old adults. mRuby signal is pseudocolored magenta. Boxed regions were magnified 5x and displayed on the right. BF, bright field. Scale bar = 10 μm. (C-F) as in (B) but represent versions of TNGL-1::mRuby fusion protein with the C-terminus progressively truncated.

**Figure S4.**
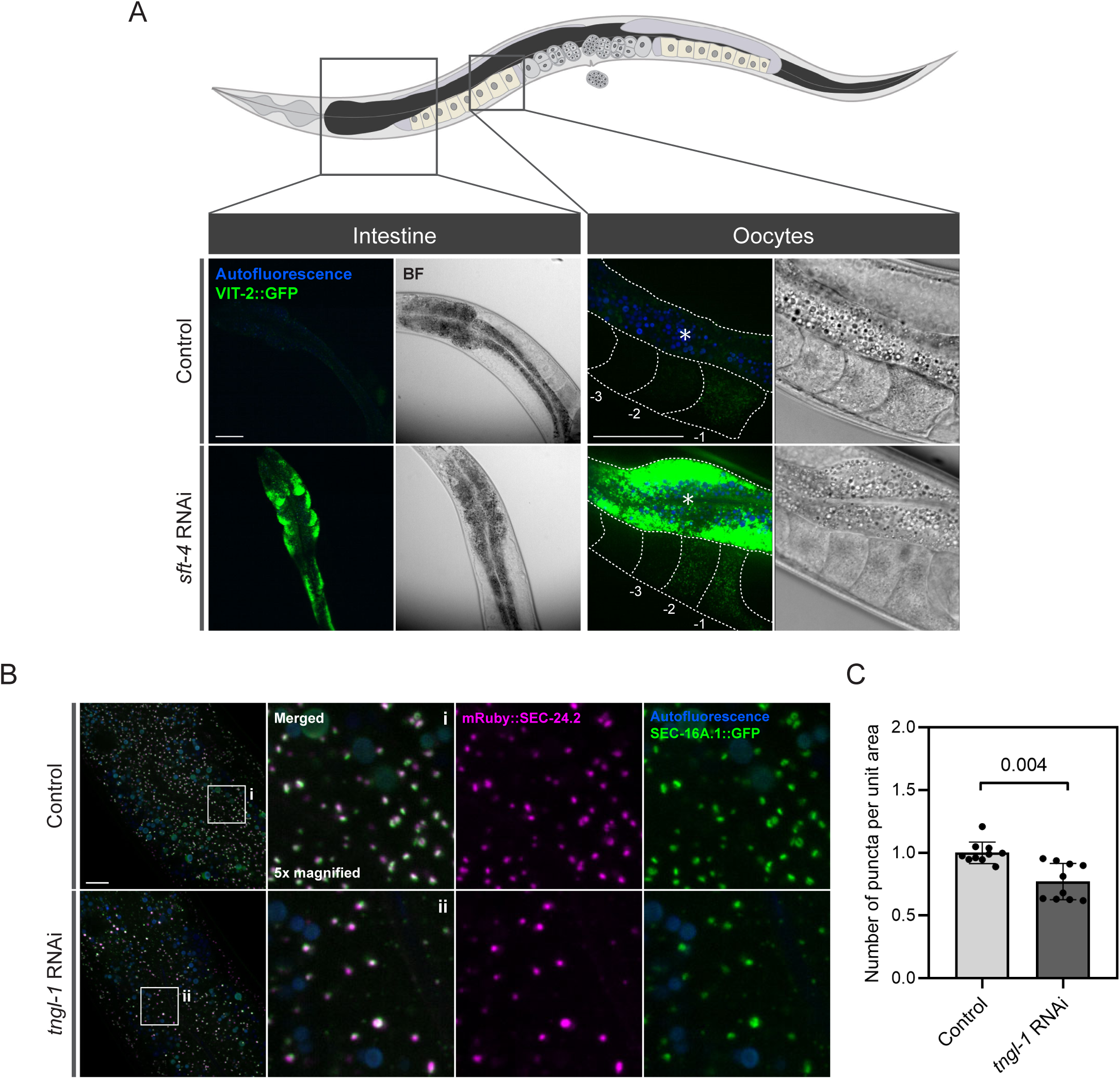
SFT-4 and TNGL-1 have distinct roles in ER cargo export. (A) SFT-4 depletion causes VIT-2::GFP accumulation in the intestine. Top, a simplified diagram of an adult-stage *C. elegans*. Images of 1-day-old adult animals are displayed. The intestine is shown in dark grey and oocytes in oocytes pastel yellow. Bottom, visualization of VIT-2::GFP (*crg9070*) in the intestine and oocytes upon treatment with control or *sft-4* RNAi. Images are projections of a 1 μm z stack. For oocytes images, the most proximal oocytes were captured. Dotted lines mark the boundary between different tissues or oocytes. Asterisks mark the intestine. The oocytes closest to the spermatheca are denoted as –1, –2 and –3 respectively. BF, bright field. Scale bars = 40 μm. (B) *tngl-1* RNAi causes aggregation of ER exit sites. ER exit sites in 1-day-old adults are visualized using mRuby::SEC24.2 (*hj331*) and SEC-16A.1::GFP (*hj256*). mRuby signal is pseudocolored magenta. Images are projections of a 1 μm z stack reconstituted from 3 focal planes. Scale bar = 10 μm. (C) Quantification of the number of SEC-16A.1::GFP positive puncta per unit area in control or *tngl-1* RNAi treated animals. n = 10 for each group. The *p*-values are displayed (unpaired t-test). Mean ± SD is shown.

## References

1. Baker, M.E. (1988). Is vitellogenin an ancestor of apolipoprotein B-100 of human low-density lipoprotein and human lipoprotein lipase? Biochem J 255, 1057–1060. 10.1042/bj2551057.

2. Babin, P.J., Bogerd, J., Kooiman, F.P., Van Marrewijk, W.J., and Van der Horst, D.J. (1999). Apolipophorin II/I, apolipoprotein B, vitellogenin, and microsomal triglyceride transfer protein genes are derived from a common ancestor. J Mol Evol 49, 150–160. 10.1007/pl00006528.

3. Smolenaars, M.M.W., Madsen, O., Rodenburg, K.W., and Van der Horst, D.J. (2007). Molecular diversity and evolution of the large lipid transfer protein superfamily. J Lipid Res 48, 489–502. 10.1194/jlr.R600028-JLR200.

4. Fisher, E.A., and Ginsberg, H.N. (2002). Complexity in the secretory pathway: the assembly and secretion of apolipoprotein B-containing lipoproteins. J Biol Chem 277, 17377–17380. 10.1074/jbc.R100068200.

5. Tiwari, S., and Siddiqi, S.A. (2012). Intracellular trafficking and secretion of VLDL. Arterioscler Thromb Vasc Biol 32, 1079–1086. 10.1161/ATVBAHA.111.241471.

6. Wahli, W. (1988). Evolution and expression of vitellogenin genes. Trends Genet 4, 227–232. 10.1016/0168-9525(88)90155-2.

7. Tufail, M., Nagaba, Y., Elgendy, A.M., and Takeda, M. (2014). Regulation of vitellogenin genes in insects. Entomological Science 17, 269–282. 10.1111/ens.12086.

8. Perez, M.F., and Lehner, B. (2019). Vitellogenins – Yolk Gene Function and Regulation in Caenorhabditis elegans. Front Physiol 10, 1067. 10.3389/fphys.2019.01067.

9. Kimble, J., and Sharrock, W.J. (1983). Tissue-specific synthesis of yolk proteins in Caenorhabditis elegans. Dev Biol 96, 189–196. 10.1016/0012-1606(83)90322-6.

10. MacMorris, M., Broverman, S., Greenspoon, S., Lea, K., Madej, C., Blumenthal, T., and Spieth, J. (1992). Regulation of vitellogenin gene expression in transgenic Caenorhabditis elegans: short sequences required for activation of the vit-2 promoter. Mol Cell Biol 12, 1652–1662. 10.1128/mcb.12.4.1652-1662.1992.

11. Yi, W., and Zarkower, D. (1999). Similarity of DNA binding and transcriptional regulation by Caenorhabditis elegans MAB-3 and Drosophila melanogaster DSX suggests conservation of sex determining mechanisms. Development 126, 873– 881. 10.1242/dev.126.5.873.

12. Hall, D.H., Winfrey, V.P., Blaeuer, G., Hoffman, L.H., Furuta, T., Rose, K.L., Hobert, O., and Greenstein, D. (1999). Ultrastructural features of the adult hermaphrodite gonad of Caenorhabditis elegans: relations between the germ line and soma. Dev Biol 212, 101–123. 10.1006/dbio.1999.9356.

13. Grant, B., and Hirsh, D. (1999). Receptor-mediated endocytosis in the Caenorhabditis elegans oocyte. Mol Biol Cell 10, 4311–4326. 10.1091/mbc.10.12.4311.

14. Ruf, H., and Gould, B.J. (1999). Size distributions of chylomicrons from human lymph from dynamic light scattering measurements. Eur Biophys J 28, 1–11. 10.1007/s002490050178.

15. Pitman, J.L., Bonnet, D.J., Curtiss, L.K., and Gekakis, N. (2011). Reduced cholesterol and triglycerides in mice with a mutation in Mia2, a liver protein that localizes to ER exit sites. J Lipid Res 52, 1775–1786. 10.1194/jlr.M017277.

16. Santos, A.J.M., Nogueira, C., Ortega-Bellido, M., and Malhotra, V. (2016). TANGO1 and Mia2/cTAGE5 (TALI) cooperate to export bulky pre-chylomicrons/VLDLs from the endoplasmic reticulum. J Cell Biol 213, 343–354. 10.1083/jcb.201603072.

17. Wang, Y., Liu, L., Zhang, H., Fan, J., Zhang, F., Yu, M., Shi, L., Yang, L., Lam, S.M., Wang, H., et al. (2016). Mea6 controls VLDL transport through the coordinated regulation of COPII assembly. Cell Res 26, 787–804. 10.1038/cr.2016.75.

18. Lekszas, C., Foresti, O., Raote, I., Liedtke, D., König, E.-M., Nanda, I., Vona, B., De Coster, P., Cauwels, R., Malhotra, V., et al. (2020). Biallelic TANGO1 mutations cause a novel syndromal disease due to hampered cellular collagen secretion. Elife 9, e51319. 10.7554/eLife.51319.

19. Guillemyn, B., Nampoothiri, S., Syx, D., Malfait, F., and Symoens, S. (2021). Loss of TANGO1 Leads to Absence of Bone Mineralization. JBMR Plus 5, e10451. 10.1002/jbm4.10451.

20. Saito, K., Chen, M., Bard, F., Chen, S., Zhou, H., Woodley, D., Polischuk, R., Schekman, R., and Malhotra, V. (2009). TANGO1 facilitates cargo loading at endoplasmic reticulum exit sites. Cell 136, 891–902. 10.1016/j.cell.2008.12.025.

21. Raote, I., Ortega-Bellido, M., Santos, A.J., Foresti, O., Zhang, C., Garcia-Parajo, M.F., Campelo, F., and Malhotra, V. (2018). TANGO1 builds a machine for collagen export by recruiting and spatially organizing COPII, tethers and membranes. Elife 7, e32723. 10.7554/eLife.32723.

22. Santos, A.J.M., Raote, I., Scarpa, M., Brouwers, N., and Malhotra, V. (2015). TANGO1 recruits ERGIC membranes to the endoplasmic reticulum for procollagen export. Elife 4, e10982. 10.7554/eLife.10982.

23. Raote, I., Ernst, A.M., Campelo, F., Rothman, J.E., Pincet, F., and Malhotra, V. (2020). TANGO1 membrane helices create a lipid diffusion barrier at curved membranes. Elife 9, e57822. 10.7554/eLife.57822.

24. Raote, I., Saxena, S., Campelo, F., and Malhotra, V. (2021). TANGO1 marshals the early secretory pathway for cargo export. Biochim Biophys Acta Biomembr 1863, 183700. 10.1016/j.bbamem.2021.183700.

25. Raote, I., and Malhotra, V. (2021). Tunnels for Protein Export from the Endoplasmic Reticulum. Annu Rev Biochem 90, 605–630. 10.1146/annurev-biochem-080120-022017.

26. Camacho, C., Coulouris, G., Avagyan, V., Ma, N., Papadopoulos, J., Bealer, K., and Madden, T.L. (2009). BLAST+: architecture and applications. BMC Bioinformatics 10, 421. 10.1186/1471-2105-10-421.

27. Söding, J., Biegert, A., and Lupas, A.N. (2005). The HHpred interactive server for protein homology detection and structure prediction. Nucleic Acids Res 33, W244–248. 10.1093/nar/gki408.

28. Gabler, F., Nam, S.-Z., Till, S., Mirdita, M., Steinegger, M., Söding, J., Lupas, A.N., and Alva, V. (2020). Protein Sequence Analysis Using the MPI Bioinformatics Toolkit. Curr Protoc Bioinformatics 72, e108. 10.1002/cpbi.108.

29. Jumper, J., Evans, R., Pritzel, A., Green, T., Figurnov, M., Ronneberger, O., Tunyasuvunakool, K., Bates, R., Žídek, A., Potapenko, A., et al. (2021). Highly accurate protein structure prediction with AlphaFold. Nature 596, 583–589. 10.1038/s41586-021-03819-2.

30. Ma, W., and Goldberg, J. (2016). TANGO1/cTAGE5 receptor as a polyvalent template for assembly of large COPII coats. Proc Natl Acad Sci U S A 113, 10061– 10066. 10.1073/pnas.1605916113.

31. Xu, N., Zhang, S.O., Cole, R.A., McKinney, S.A., Guo, F., Haas, J.T., Bobba, S., Farese, R.V., and Mak, H.Y. (2012). The FATP1-DGAT2 complex facilitates lipid droplet expansion at the ER-lipid droplet interface. J Cell Biol 198, 895–911. 10.1083/jcb.201201139.

32. Wang, X., Wang, H., Xu, B., Huang, D., Nie, C., Pu, L., Zajac, G.J.M., Yan, H., Zhao, J., Shi, F., et al. (2021). Receptor-Mediated ER Export of Lipoproteins Controls Lipid Homeostasis in Mice and Humans. Cell Metab 33, 350–366.e7. 10.1016/j.cmet.2020.10.020.

33. Saegusa, K., Sato, M., Morooka, N., Hara, T., and Sato, K. (2018). SFT-4/Surf4 control ER export of soluble cargo proteins and participate in ER exit site organization. J Cell Biol 217, 2073–2085. 10.1083/jcb.201708115.

34. Tufail, M., and Takeda, M. (2008). Molecular characteristics of insect vitellogenins. J Insect Physiol 54, 1447–1458. 10.1016/j.jinsphys.2008.08.007.

35. Gottlieb, T.A., and Wallace, R.A. (1982). Intracellular glycosylation of vitellogenin in the liver of estrogen-stimulated Xenopus laevis. J Biol Chem 257, 95–103.

36. Sharrock, W.J., Sutherlin, M.E., Leske, K., Cheng, T.K., and Kim, T.Y. (1990). Two distinct yolk lipoprotein complexes from Caenorhabditis elegans. J Biol Chem 265, 14422–14431.

37. Roy-Bellavance, C., Grants, J.M., Miard, S., Lee, K., Rondeau, É., Guillemette, C., Simard, M.J., Taubert, S., and Picard, F. (2017). The R148.3 Gene Modulates Caenorhabditis elegans Lifespan and Fat Metabolism. G3 (Bethesda) 7, 2739–2747. 10.1534/g3.117.041681.

38. Sirwi, A., and Hussain, M.M. (2018). Lipid transfer proteins in the assembly of apoB-containing lipoproteins. J Lipid Res 59, 1094–1102. 10.1194/jlr.R083451.

39. Chatterton, J.E., Phillips, M.L., Curtiss, L.K., Milne, R., Fruchart, J.C., and Schumaker, V.N. (1995). Immunoelectron microscopy of low density lipoproteins yields a ribbon and bow model for the conformation of apolipoprotein B on the lipoprotein surface. J Lipid Res 36, 2027–2037.

40. Bouchoux, J., Beilstein, F., Pauquai, T., Guerrera, I.C., Chateau, D., Ly, N., Alqub, M., Klein, C., Chambaz, J., Rousset, M., et al. (2011). The proteome of cytosolic lipid droplets isolated from differentiated Caco-2/TC7 enterocytes reveals cell-specific characteristics. Biol Cell 103, 499–517. 10.1042/BC20110024.

41. Berndsen, Z.T., and Cassidy, C.K. (2024). The Structure of ApoB100 from Human Low-density Lipoprotein. Preprint, 10.1101/2024.02.28.582555 https://doi.org/10.1101/2024.02.28.582555.

42. Zhai, C., Zhang, N., Li, X.-X., Tan, X.-K., Sun, F., and Dong, M.-Q. (2024). Immuno-electron microscopy localizes *Caenorhabditis elegans* vitellogenins along the classic exocytosis route. Life Metabolism, loae025. 10.1093/lifemeta/loae025.

43. Zhang, Z., Bai, M., Barbosa, G.O., Chen, A., Wei, Y., Luo, S., Wang, X., Wang, B., Tsukui, T., Li, H., et al. (2020). Broadly conserved roles of TMEM131 family proteins in intracellular collagen assembly and secretory cargo trafficking. Sci Adv 6, eaay7667. 10.1126/sciadv.aay7667.

44. Zhang, Z., Luo, S., Barbosa, G.O., Bai, M., Kornberg, T.B., and Ma, D.K. (2021). The conserved transmembrane protein TMEM-39 coordinates with COPII to promote collagen secretion and regulate ER stress response. PLoS Genet 17, e1009317. 10.1371/journal.pgen.1009317.

45. Liu, M., Feng, Z., Ke, H., Liu, Y., Sun, T., Dai, J., Cui, W., and Pastor-Pareja, J.C. (2017). Tango1 spatially organizes ER exit sites to control ER export. J Cell Biol 216, 1035–1049. 10.1083/jcb.201611088.

46. Neve, I.A.A., Sowa, J.N., Lin, C.-C.J., Sivaramakrishnan, P., Herman, C., Ye, Y., Han, L., and Wang, M.C. (2020). Escherichia coli Metabolite Profiling Leads to the Development of an RNA Interference Strain for Caenorhabditis elegans. G3 (Bethesda) 10, 189–198. 10.1534/g3.119.400741.

47. Klemm, R.W., Norton, J.P., Cole, R.A., Li, C.S., Park, S.H., Crane, M.M., Li, L., Jin, D., Boye-Doe, A., Liu, T.Y., et al. (2013). A conserved role for atlastin GTPases in regulating lipid droplet size. Cell Rep 3, 1465–1475. 10.1016/j.celrep.2013.04.015.

48. Cao, Z., Fung, C.W., and Mak, H.Y. (2022). A Flexible Network of Lipid Droplet Associated Proteins Support Embryonic Integrity of C. elegans. Front Cell Dev Biol 10, 856474. 10.3389/fcell.2022.856474.

49. Brenner, S. (1974). The genetics of Caenorhabditis elegans. Genetics 77, 71–94. 10.1093/genetics/77.1.71.

50. Dickinson, D.J., Pani, A.M., Heppert, J.K., Higgins, C.D., and Goldstein, B. (2015). Streamlined Genome Engineering with a Self-Excising Drug Selection Cassette. Genetics 200, 1035–1049. 10.1534/genetics.115.178335.

51. Labun, K., Montague, T.G., Krause, M., Torres Cleuren, Y.N., Tjeldnes, H., and Valen, E. (2019). CHOPCHOP v3: expanding the CRISPR web toolbox beyond genome editing. Nucleic Acids Res 47, W171–W174. 10.1093/nar/gkz365.

52. Frøkjær-Jensen, C., Davis, M.W., Ailion, M., and Jorgensen, E.M. (2012). Improved Mos1-mediated transgenesis in C. elegans. Nat Methods 9, 117–118. 10.1038/nmeth.1865.

